# Genomic Regions and Variant Analysis Reveal Candidate Genes Associated with Age at First Service in Chinese Holstein Heifers

**DOI:** 10.1101/2025.07.28.667120

**Authors:** Cuili Pan, Longgang Ma, Lulu Wang, Zhuangbiao Zhang, Ao Wang, Fengting Bai, Yudong Cai, Yun Ma, Yachun Wang, Guanglei Liu, Kai Zhu, Xinzhe Lv, Xihong Wang, Yu Jiang

## Abstract

Understanding the genetic basis and identifying quantitative trait loci (QTL) for age at first service (AFS) is essential for improving reproductive efficiency and reducing economic costs in heifers rearing. We conducted a genome-wide association study (GWAS) for AFS using the genomic estimated breeding values (GEBVs) of 3,686 Chinese Holstein heifers, genotyped with 7,964,217 single nucleotide polymorphisms (SNPs) after quality control. Significant SNPs and candidate genes were identified and investigated through colocalization analysis of GWAS, expression QTL (eQTL), and splicing QTL (sQTL), and expression analysis in blood RNA sequencing (RNA-seq). The heritability estimate for AFS was moderate at 0.276 ±0.025. Three QTL regions associated with AFS were identified: Region 1 on BTA6 (43688837-45007127 bp, 1.3 Mb with three 200kb windows), explaining 0.47% of genetic variance; Region 2 on BTA6 (62770788-62967316 bp) explaining 0.27% of the genetic variance; and Region 3 on BTA18 (6299721-6498934 bp) explaining 0.17% of the genetic variance. Combining GWAS and colocalization analysis, we identified *ANAPC4, RBPJ, SEPSECS, SLC34A2, ZCCHC4*, CCDC149, *GNPDA2, GUF1, DHX15, SOD3, GUF1* and *GNPDA2* as candidate genes. These genes are enriched in signaling pathways such as progesterone-mediated oocyte maturation, parathyroid hormone synthesis, secretion and action, and selenocompound metabolism. *ANAPC4, DHX15, GUF1, SEPSECS*, and *SOD3* were also differently expressed genes (DEGs) in blood RNA-seq. Integrating the GWAS, colocalization, differential expression analysis, and existing literature, *ANAPC4* and rs136363104 were highlighted as the putative causal gene and variant for puberty, respectively. In conclusion, our study identified three candidate regions and 12 candidate genes, confirming *ANAPC4* as a putative causal gene influencing AFS. This research enhances the understanding of the genetic basis of AFS in Chinese Holstein heifers. The identified key genomic regions, candidate genes, and variants have the potential to improve reproductive efficiency and reduce economic costs in heifer rearing.

**Interpretative summary:** The economic burden of heifer-raising and reproductive inefficiency in dairy cattle is evident. The most important strategy for optimizing heifer-raising is selecting heifers with earlier puberty (sexual maturity) to shorten age at first service (AFS). In this study, we have identified key genomic regions and candidate genes on AFS in Chinese Holstein heifers by genome-wide association study, and validated them by colocalization and RNA-seq analysis. The findings have the potential to enhance the understanding of the genetic basis of reproductive traits, leading to more targeted breeding strategies for improved reproductive efficiency.

## INTRODUCTION

Age at first service (AFS) is an economically important trait as it determines the timepoint a heifer starts its reproductive life and affects age at first calving (AFC), which impacts generation interval and rearing costs. Although heifers are the future revenue-generating units of a herd, the nonproductive rearing period (RP) is a large investment without financial return until heifers enter the milking period, which inevitably brings great economic pressure for the dairy farms. The average rearing costs was $2355 in total and $2.45 per day per heifer according to the estimations using the inputs and investments information of 26 farms in the United States, which might account for as much as 15% to 20% of total operating expenses (Karszes et al., 2020). Therefore, any strategy that aimed to reduce the cost of the heifer-raising might have a pronounced influence on the overall profitability of dairy industry.

Of particular interest are the genetic improvement strategies to reduce the AFC, which shorten the time to incorporate heifers into the milking herd, thereby achieving a positive lifetime cash flow earlier (Ettema et al., 2004). Among the fertility traits, AFC is composed of AFS, the time from artificial insemination to pregnancy (IFLh), and the gestation length (GL) (Heise et al., 2018). GL is relatively fixed, averaging 273 days in Chinese Holstein heifers in this study and 277.8 days previously reported in US Holstein heifers (Norman et al., 2009). IFLh is the time from first to last (or successful) insemination, depending on the sexual maturity and ability to conceive of the heifers (Weller et al., 2022a). Therefore, the most important strategy for optimizing AFC is selecting heifers with earlier puberty (sexual maturity) to shorten AFS. So far, proactive genetic and genomic evaluation has been implemented, aiming to reduce the AFS and improve the insemination and conception probability after heifers reached the appropriate size to become pregnant (Kgari et al., 2022a; Prakapenka et al., 2023; Stephen et al., 2023; Ferrari et al., 2024).

As most fertility traits in dairy cattle are controlled by multiple genes, Genome-Wide Association Studies (GWAS) are widely used to explore genomic regions that contribute to genetic variation. In both beef and dairy cattle, AFS is a key fertility trait associated with productivity. Identifying mutations underpinning variation in AFS between individuals has been challenging due to large environmental variation. Notwithstanding, progress has been made, and several genomic regions and candidate genes for AFC, AFS, and age at puberty (AGEP) have been identified in cattle (Weller et al., 2022b; Prakapenka et al., 2023; Stephen et al., 2023). However, the overlap of QTL locations among different populations is generally poor, demonstrating the challenges in finding candidate genes and mapping causal variants for AFS. So far, study on the candidate regions and genes of AFS or AGEP in Chinese Holstein heifers was limited.

Benefiting from the extensive use of automated activity monitors, large amounts of information regarding the detection and behavioral profile of estrus have become available and powerful in cows (Cerri et al., 2021). Therefore, based on the collection of AFS by intelligent equipment detection, the main objectives of this study were to: 1) explore the phenotypic rules and genetic parameters of 7 fertility traits of Chinese Holstein heifers; 2) identify genetic regions, variants, and candidate genes affecting AFS by GWAS and locate important QTL/eQTL/sQTL regions; 3) study the expression of candidate genes based on RNA sequencing (RNA-seq). The results would provide clues to understand the regulation and genetic architecture of AFS in Chinese Holstein heifers.

## MATERIALS AND METHODS

### Phenotype and Genotype Data Collection and Preprocessing

In this study, no animal handling was conducted. The phenotype and genotype data were gathered from the routine breeding and genomics evaluations. Therefore, this study did not require approval by an Institutional Animal Care and Use Committee.

The birth, mating and calving records of Chinese Holstein heifers were collected between 2015 and 2024. From these records, seven fertility traits of heifers were obtained: age at first service (AFS), age at first calving (AFC), the interval from first to last insemination in heifer (IFLh), conception rate of first insemination (CRh), The number of repeated service (NS), gestation length (GL), birth weight (BiW). Phenotype data were processed using the principle of mean ± 3σ (σ is standard deviation) to remove the outliers.

Blood samples were collected and DNA was extracted for low-coverage sequencing (lcWGS) (average depth of 1×) at MolBreeding Biotech Co. Ltd (Shijiazhuang, Hebei province, China). The raw lcWGS data were subjected to quality control using fastp v0.20.0 (Chen et al., 2018) with default parameters. Subsequently, BWA MEMv0.7.17 (Li et al., 2010) was used to align the clean reads to the bovine reference genome ARS-UCD1.2. The bam file conversion, sorting, and duplicate read filtering were conducted using samtools v1.10 (Li et al., 2009) and Picard v2.20.1 (http://broadinstitute.github.io/picard/). Using the reference panel and workflow from a previous study (Zhang et al., 2023), whole-genome imputation was performed, resulting in a final set of 11,106,737 SNPs. To meet the analysis requirements, further quality control was conducted to exclude SNPs with a minor allele frequency <0.05, genotype missing rate > 0.1, sample missing rate > 0.1, and Hardy-Weinbery P-value < 1e-06. After quality control, 7,964,217 SNPs with a density distribution displayed in Fig. S1, Additional file 2 and 3,683 records of phenotype (3,683 herfers) were used for further analyses.

### Genetic Parameter Estimation

The phenotypic and genetic variances of fertility traits in Chinese Holstein heifers were estimated using the Average Information Restricted Maximum Likelihood (AI-REML) method. This estimation was performed with single linear animal models implemented through the BLUPF90+ family of programs (RENUMF90, AIREMLF90 and BLUPF90) (Misztal et al., 2014). Fixed effects of the seven fertility traits included farm, year-season of birth, mating and calving, sex of calf, gestation length (month), and first mating month. For the variant management levels and technology, samples from the farms was treated as three level in fixed effect. The birth, mating and calving years each included 6 levels (2015-2020, 2017-2021, one level per year). The twelve months were divided into four levels according to the season (spring: March to May; summer: June to August; autumn: September to November; winter: December to February). The general liner model (GLM) procedure was utilized to test the significance of the above fixed effects for each phenotype and determine the ones to be incorporated in the genetic evaluation.

The single-trait animal model was used to estimate (co)variance components:

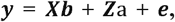

where **y** is the vector of phenotypic records for AFS, AFC, IFLh, NSh, CRh, GLh, BiWh; **b** is the vector of fixed/systematic effects (farm and year-season of birth for AFS, farm and year-season of mating for AFC and IFLh, farm and year-season of calving for GLh, year-season of mating and the first mating month for NSh, farm, year-season of calving, sex of calf and gestation length for BiWh); ***a*** is a vector of random additive genetic effects; 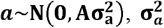 is the additive genetic variance, **A** is the genotype-based relationship matrix; and **e** is the vector of random residual effects; 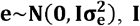 is and identity matrix, and 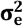 is the residual variance; **X, Y** are incidence matrices linking the corresponding phenotypic records to **b** and **a**, respectively.

The calculation formula of heritability is:

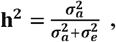

where **h**^**2**^ is the heritability, 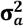,is the additive genetic variance, and 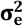 is the residual variance. The square of the SE^2^ for the heritability estimates was calculated as:

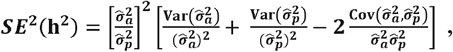

The binary-trait animal model was used to estimate genetic correlations:

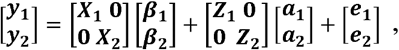

where y_1_ and y_2_ are the matrix of phenotypes; β_1_ and β_2_ are the fixed effect vector of the corresponding phenotype; ***a***_1_ and ***a***_2_ are random additive genetic effect vectors of the corresponding phenotype; e_1_ and e_2_ are the random residual effect vector for the first and second phenotypes; X, Z are incidence matrices linking the corresponding phenotypic records to β and a, respectively.

The genetic correlation coefficient was calculated as 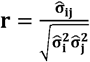, and the square of the SE for the genetic correlation coefficient was calculated as(Su et al., 2007):

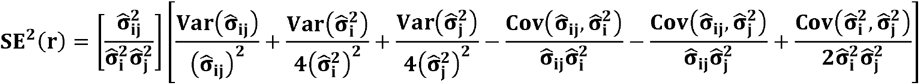

The variances and covariances of the estimated parameters 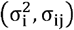 were obtained from the inverse of the average information matrix.

### Genome-Wide Association Study

#### Exploration of Genomic Regions

The POSTGSF90 module in BLUPF90 family was used to explore the genetic variation explaination ratio of AFS in the genome-wide regions. Genome-wide linkage disequilibrium (LD) attenuation analysis determined 200kb as the maximum linkage distance and used it as the window size in further study. The genomic best linear unbiased prediction (BLUP) of AFSs was analyzed with a single trait animal model described above, and the percentage of additive genetic variance explained by the *i-*th SNP window was calculated as:

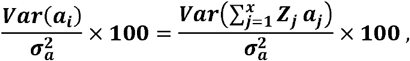

where ***a***_***i***_ is the genetic value of the *i-*th genomic region, 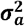 is the total additive genetic variance, ***x*** is the total number of adjacent SNPs within the 200 kb region, ***Z***_***j***_ is the vector of gene content of the *j-*th SNP for all windows, and ***a***_***j***_ is the effect of the *j-*th SNP within *i-*th window. The results are reported as the percentage of genetic additive variance explained by 200 kb windows and plotted using ggplot2 package in R software (version 4.3.0). Additionally, we calculated the P-value of individual SNP for GWAS from elements of the inverse of the Mixed Model Equations previously obtained from blupf90 (Aguilar et al., 2019).

#### Identification of Significant SNP

The linear mixed model (LMMs, also known as mixed linear models, or MLMs) was used to test the association in our studies. This method corrects population structure by taking into account the relatedness matrix, which is built by considering all genome-wide SNPs as a random effect. The statistical model for single marker regression analysis can be described as:

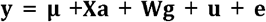

where **y** is the vector of GEBVs of AFSs for n individuals; **μ** is the overall mean effect (as same as intercept term); X is an matrix of fixed effects as same as ssGWAS; **a** is a c-vector of the corresponding fixed effects including the intercept; W is an n-vector of marker genotypes, **g** is the effect size of the SNP maker (allele substitution effect); u is the n-vector of random polygenic effects, and e is an n-vector of errors. The vector of random effects was assumed to follow an n-dimensional multivariate normal (MVN) distribution u ∼ MVN_n_,(0,λ τ^−1^K), where τ^−1^ is the variance of residual errors, *λ* is the ratio between the two variance components, and K is a known n × n identity matrix.

The genomic inflation factor (λ) was the ratio of the statistic median of the observed distribution to the expected, which could be calculated by the formula: *λ* = median,(*χ*^2^)/0.4549. As a large number of statistical tests were performed, the traditional Bonferroni correction would be highly conservative because not all tests are independent due to LD. To avoid excessive false-negative results, a modified Bonferroni correction was applied, which used the number of independent chromosome segments (M_e_) instead of the total number of SNPs. M_e_ was calculated as (Goddard et al., 2011):

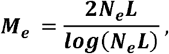

where *M*_*e*_ is a function of effective population size (*N*_*e*_) and the average chromosome length in Morgan (*L*). The most conservative value of *N*_*e*_ in Chinese Holstein cattle population was 33 (Shi et al., 2020). So in our study it was also set it as 33. The chromosome-wide statistically significant threshold was calculated by dividing 0.01 or 0.05 by *M*_*e*_ (Table S1, Additional file 1). The corresponding significance thresholds for different chromosomes ranged from 4.75/4.25 (BTA25) to 5.45/4.95 (BTA1). The significant SNPs were selected with a threshold of -log10 (P-value) higher than the chromosome-wide threshold. Using a average *M*_*e*_ across each chromosome, the genome wide significance thresholds was about 5.0. Previous study reported P < 1e-05 (−log10 (P-value) > 5) was commonly used as the alternative significance thresholds for association analyses (Consortium, 2007; Yang et al., 2010). Given the LD between SNPs and the discovery nature of this study, we chose 5.0 as the correction threshold.

#### Gene based GWAS analysis

Gene-based association analysis was further conducted using MAGMA (version 1.07) (de Leeuw et al., 2015). The cattle gene location file was generated by combining the locations of genome-wide SNPs and the bovine reference genome annotation ARS-UCD1.2. Gene analysis was performed on GWAS data in binary PLINK format (.bed/.bim/.fam files) using the previously generated annotation file. We employed the top SNP (snp-wise=top module) model with an adaptive permutation procedure to calculate the association of each gene with AFS, which is particularly sensitive when only a small percentage of SNPs in a gene are correlated with the trait. To include SNPs in a window around genes, the window modifier was set as ‘--annotate window=5,5’, specifying SNPs within a 5kb upstream and downstream window for analysis. Bonferroni correction was applied to find the genome-wide significant genes with P-value < 2.53e-06 (i.e., nominal type I error of 0.05 with a total number of 19,756 gene tests).

### Gene Annotation, QTL Identification and Functional Enrichment

Single SNP with the smallest P-values might not explain a large percentage of genetic variance. Therefore, we combined the percentage of variance explained and significant SNPs within consecutive 200 kb windows to identify candidate genomic regions for further investigation. Genes within the five windows explaining the largest amount of variance were annotated using the ANNOVAR software (Wang et al., 2010; Yang et al., 2015) (http://annovar.openbioinformatics.org/) with the *Bos taurus* ARS-UCD 1.2 genome assembly (http://bovinegenome.org). Functional annotation of the identified genes was performed using the clusterProfiler R package (Yu et al., 2012; Wu et al., 2021) and the Kyoto Encyclopedia of Genes and Genomes (KEGG) pathway database (https://www.genome.jp/kegg). The annotation and enrichment analyses of QTLs were performed using the GALLO v.0.99 R package (Fonseca et al., 2020) and the Animal QTLdb database. Enriched QTLs were identified from the annotated QTL using the number of annoatted QTLs within the candidate regions and the total number in the database, employing a hypergeometric test with a threshold value of FDR <0.05. Cis expression quantitative trait locus (cis-eQTL) and cis splicing quantitative trait locus (cis-sQTL) were detected using fastQTL (Mohammadi et al., 2017), and trans-eQTL was detected by a mixed linear model using the parameter “-mlma” from GCTA in 7,180 publicly available RNA-Seq samples from Genotype-Tissue Expression atlas (cattleGTEx) (Ongen et al., 2017; Liu et al., 2022).

### Transcriptome Sequencing and Expression Analysis

#### Library Preparation and RNA Sequencing

A total of 196 individuals with AFS ranging from 395d to 487d were randomly selected from Holstein heifers for RNA-seq. Blood sample were collected, and total RNA was isolated using the TRIzol Reagent method according to the manufacturer’s instructions. RNA concentration, quality and integrity were determined using a Nanodrop 2000 (Thermo, Massachusetts, USA), Equalbit RNA BR Assay Kit (Cat No. Q10211, Invitrogen, Carlsbad, CA, USA), and 1% agarose gel electrophoresis. High quality RNA was used to construct cDNA library with the NEBNext Ultra RNA Library Prep Kit for Illumina (Cat No. E7530S, New England Biolabs Ltd., Hitchin, Herts, UK). Sequencing was performed on the NovaSeq 6000 System (Illumina Inc., San Diego, CA, USA), generating 150 bp paired-end reads.

#### Quality Control, Reads Alignment and expression analysis

Sequencing quality was assessed using FastQC (version 0.11.9), and raw sequence data were processed using Trim Galore (version 0.6.6) to remove adapter sequences, low-quality reads, and short reads. Clean reads were then mapped to the bovine reference genome ARS-UCD1.2 using STAR (version2.7.11a) software (Dobin et al., 2015), and the expression levels of all genes and transcripts were quantified using RESM (version 1.3.3) software (Li et al., 2011). Principal component analysis (PCA) of read counts was performed using the FactoMineR (version 2.9) R package. Differentially expressed genes (DEGs) were identified using the limma (version 3.19) package (Ritchie et al., 2015) in R, with a significance threshold of adjusted P-value <0.05 and an absolute value of fold change (|FC|) > 1.2. Gene and transcripts expression analysis, clustering analysis (k-means clustering algorithm), and pairwise comparisons were also conducted and visualized in R (version 4.3.0) software.

## RESULTS

### Descriptive Statistics and Genetic Parameters Estimation

The number of records (N), mean, standard deviation (SD), coefficient of variation (CV), median, mininum, and maxinum for seven fertility traits in Chinese Holstein heifer are shown in Table 1. The mean of AFS was 421.91±21.84 days, ranging from 367 to 560 with a variable coefficient of 5.18%. AFS, AFC and GLh had low CVs (less than 10%), while IFLh, NSh, and CRh had high CVs (186.14%, 60.95%, and 85.07%, respectively). The phenotype (AFS) slightly deviated from the normal distribution (Fig. 1A).

**Table 1.**
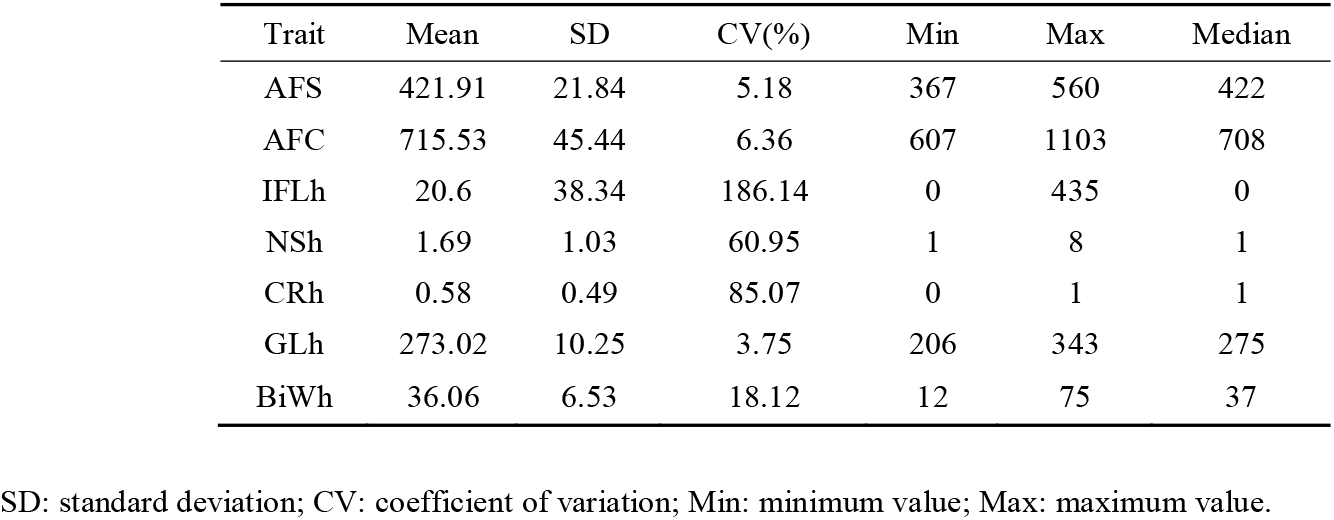
Descriptive statistics of fertility trait in Chinese Holstein heifer.

**Fig. 1.**
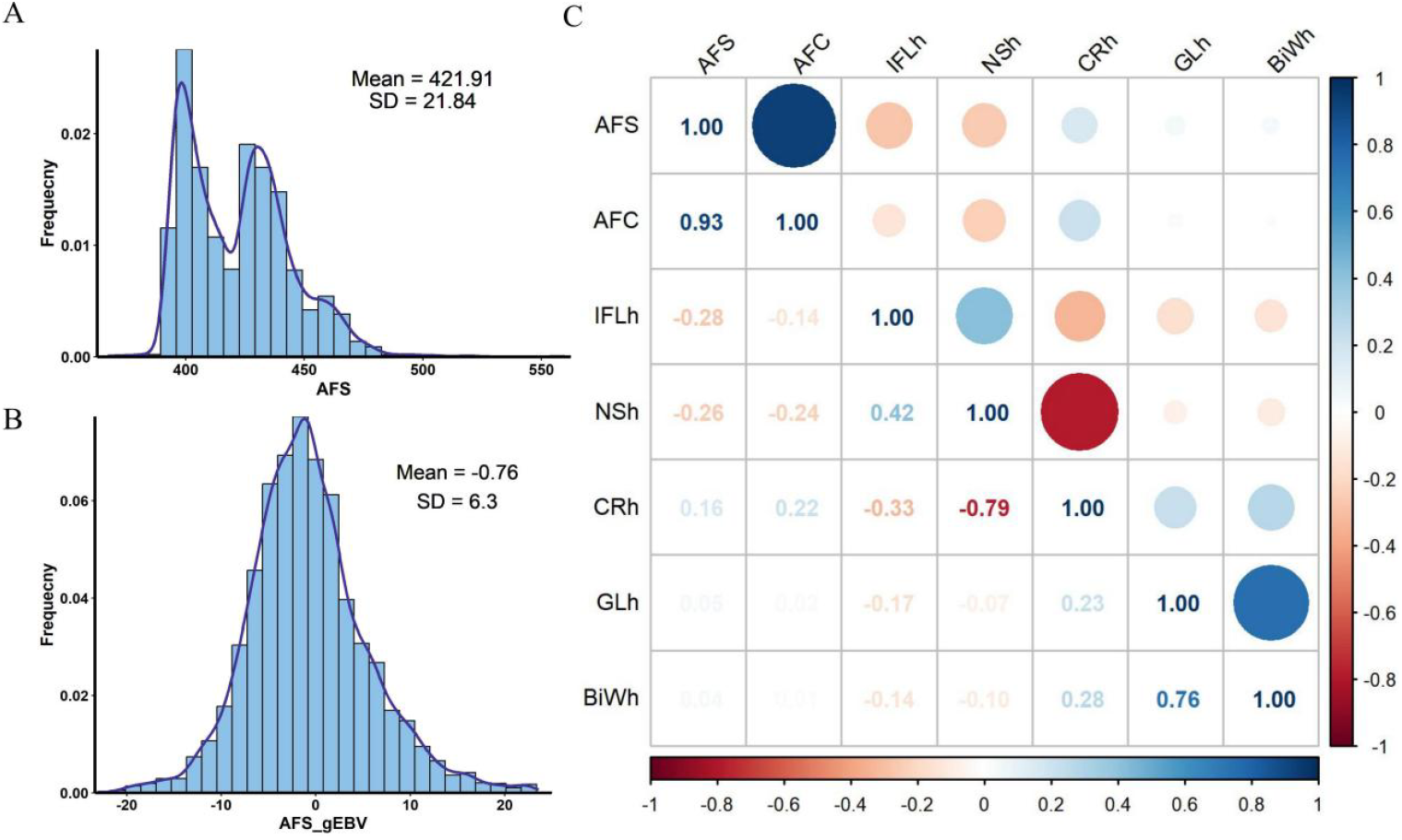
The phenotype, GEBV and genetic correlation of AFS in Chinese Holsteins heifers. **(A)** The phenotype distribution of AFS. The X-axis is the days of AFS, and the Y-axis is the frequency of individuals. (B) The genome estimated breeding value (GEBV) distribution of AFS. The X-axis is the EBV, and the Y-axis is the frequency of individuals. (C) Genetic correlations between AFS and other six fertility traits. Blue represents a positive correlation and red represents a negative correlation.

Variance component and heritability of the seven fertility traits was calculated and are displyed in Table 2. It was found that AFS had a moderate heritability of 0.276, while AFC, IFLh, NSh, CRh, GLh, and BiWh showed low heritability. The GEBV of AFS, used for the subsequent GWAS analysis, was estimated, and its distribution is shown in Fig 1B.

**Table 2.** The variance component and heritability of fertility traits in Chinese Holstein heifer.

| Trait | V<sub>a</sub> | V<sub>e</sub> | SE(Var) | h<sup>2</sup> | SE(h<sup>2</sup>) | P-value(h<sup>2</sup>) |
| --- | --- | --- | --- | --- | --- | --- |
| AFS | 71.474 | 187.762 | 7.349 | 0.276 | 0.025 | 1.89E-28 |
| AFC | 97.449 | 1604.560 | 28.870 | 0.057 | 0.017 | 6.27E-04 |
| IFLh | 1.372 | 1383.990 | 15.084 | 0.001 | 0.011 | 9.28E-02 |
| NSh | 0.035 | 0.960 | 0.016 | 0.035 | 0.016 | 2.75E-02 |
| CRh | 0.007 | 0.227 | 0.004 | 0.029 | 0.016 | 6.63E-02 |
| GLh | 4.711 | 91.306 | 1.554 | 0.049 | 0.016 | 2.20E-03 |
| BiWh | 2.736 | 18.185 | 0.516 | 0.131 | 0.024 | 3.24E-08 |

V<sub>a</sub>: variance of additive genetic effect; V<sub>e</sub>: variance of residual effect; Var: variance; SE: standard error; h<sup>2</sup>: heritability.

Genetic correlation analysis showed that AFS had a high positive correlation of 0.93 with AFC (Fig 1C), which is consistent with the understanding that early AFS can advance the production period by bringing AFC forward. Among other traits, AFS showed a moderate negative genetic correlation with IFLh and NSh (−0.08 and -0.26, respectively). There was a low positive genetic correlation between AFS and CRh (0.16). Notably, there was a moderate positive genetic correlation between NSh and IFLh (0.42) and a high negative genetic correlation between CRh and NSh (−0.79).

### Exploration of Candidate Regions Based on Additive Genetic Variance Explanation

The whole genome was divided into 200kb windows, and the additive genetic variance explaination ratio of AFS by the SNPs in each window was calculated (Fig. 2A). There were 77 windows with a genetic variation explanation rate greater than 0.1%, accounting for 11.2% of the total (Table S2, Additional file 1). The top1% windows explained 13.38% of the total additive genetic variation. Among these, 14 windows on BTA6 accounted for the most (2.03%), 9 windows on BTA9 accounted for the second (1.30%), and 6 windows on BTA18 the third (0.93%). The single window with the largest variation explanation was also located on BTA6:62770788-62967316, explaining 0.27% of the genetic variation. Additionally, three windows on BTA6 within 1.32Mb (43688837-6:45007127) together accounted for 0.47% of the genetic variation (Fig. 2C), and were subsequently analyzed as a joint region named Region 1. In summary, based on the additive effect explanation percentage of AFS, we focused on candidate regions located on BTA6, BTA9 and BTA18 for further study.

**Fig. 2.**
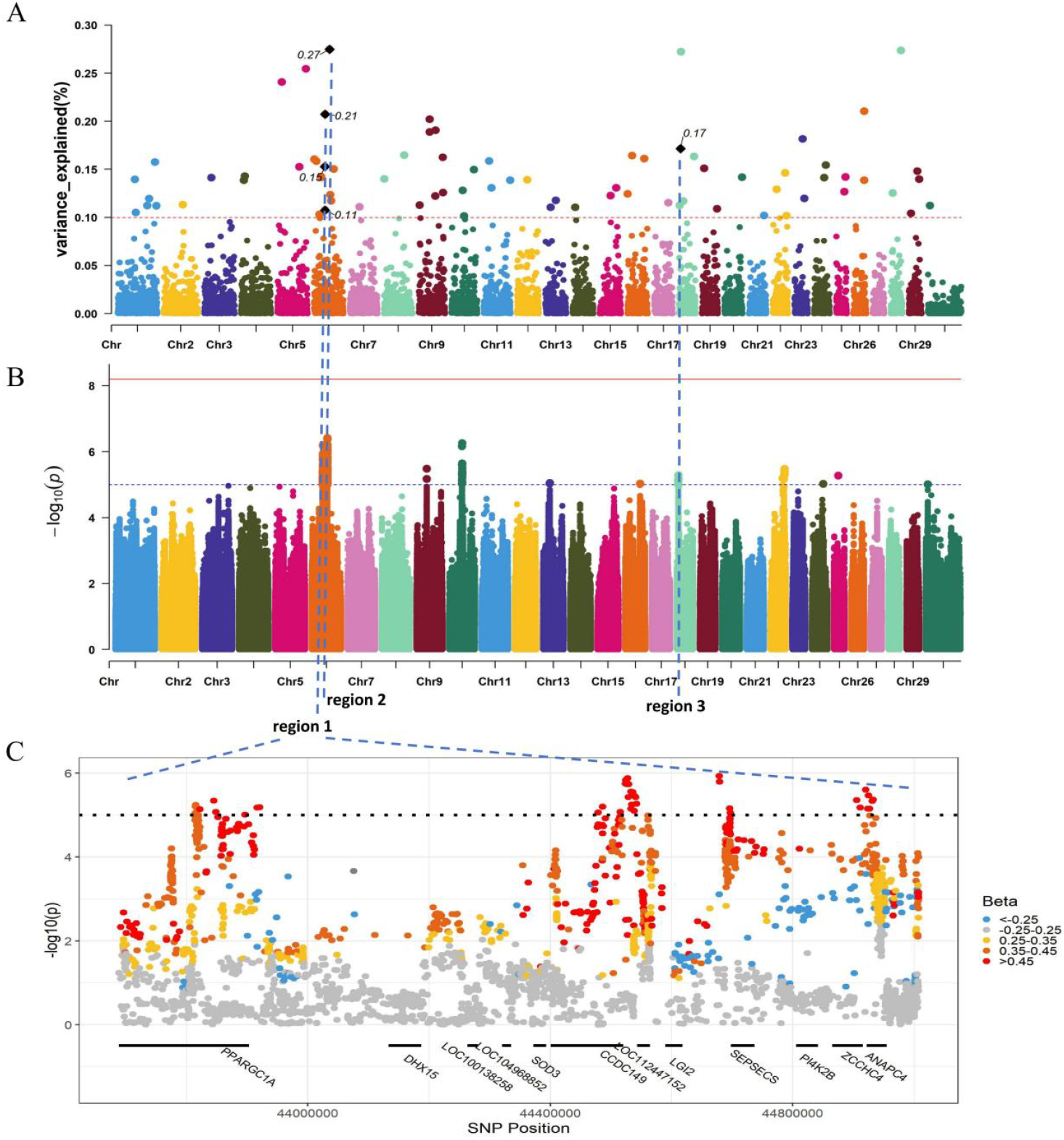
Identification of candidate regions and variants of AFS by GWAS. (A) Manhattan plot for the proportion of genetic variance explained by nonoverlapping windows of 200kb for AFS. The horizontal axis represents the chromosome and location of each window, and the vertical axis represents the percentage of variance explained. (B) Manhattan plot displaying the association of genome-wide SNPs with AFS. The vertical axis depicts –log10(p) values for the variants. The solid, horizontal red line denotes the genome-wide Bonferroni correction significance threshold at p = 6.28e-09, and the dashed, horizontal blue line denotes the modified Bonferroni correction threshold at p = 1e-05. Points above the blue line represent significant hits. (C) Manhattan plot of genome-wide p-values for the identified candidate region 1. The vertical axis represents –log10(p) values and the color depicts the effect (Beta) of each SNP. The genes within the region are shown in the bottom.

### Identification of Candidate Regions Combined with SNP Significance

We identified 10 loci significantly associated with AFS: two on BTA6 and the others on BTA9, BTA10, BTA13, BTA16, BTA18, BTA22, BTA24, and BTA25, respectively (Fig. 2B; Fig. S2, Additional file 2; Table S3, Additional file 1). The Q-Q plot and the lambda value (λ = 1.011) indicate that potential population stratification seems to have been accounted for in this study (Fig. S2, Additional file 2). By combining the top 1% 200kb windows of the additive genetic variance explaination, five intersecting windows containing significant SNPs were obtained (Fig. 2). The three adjacent windows within 1.32Mb on BTA6 were merged into Region 1 (Fig. 2C), which contained 45 significant SNPs. The single window on BTA6:62770788-6:62967316 with the largest variation explanation was named Region 2, which included 69 significant SNPs. Region 3 was located on BTA18, spanning an interval from 6299721 to 6498934, containing 12 significant SNPs.

Linear-regression and correlation analysis showed that SNP significance had a significant positive correlation with its effect (r = 0.89, p = 2.2e-16) in Region 1 (Fig. S3, Additional file 2), indicating that significant SNPs tend to have relative strong effects. Consistent with this, the obtained 45 significant SNPs all showed effects greater than 0.45 (Fig. 2C). These results also supported the screening of candidate regions and variants based on SNP effect and significance in our study.

### Functional Enrichment Analysis of Candidate QTLs and Genes

QTL annotations provide a better understanding of the frequency of each QTL class in the candidate regions. The most frequently annotated QTL class was Milk (76.05%), followed by Production (15.21%), Meat and Carcass (3.24%), Reproduction (2.27%), Health (1.94%), and Exterior (1.29%) (Fig. S4, Additional file 2). We also investigated the detailed composition of the Milk, Production, and Reproduction QTLs class in our candidate regions (Fig. S5, Additional file 2). The investigation bias for milk related traits in the QTL database might result in a larger proportion of records. To reduce the bias, a QTL enrichment analysis was also performed, showing enrichment in traits related to milk (such as the percentage or content of milk protein, fat, potassium, palmitoleic acid, and casein), and production (body weight, bone weight, and body length) (Fig. 3A).

**Fig. 3.**
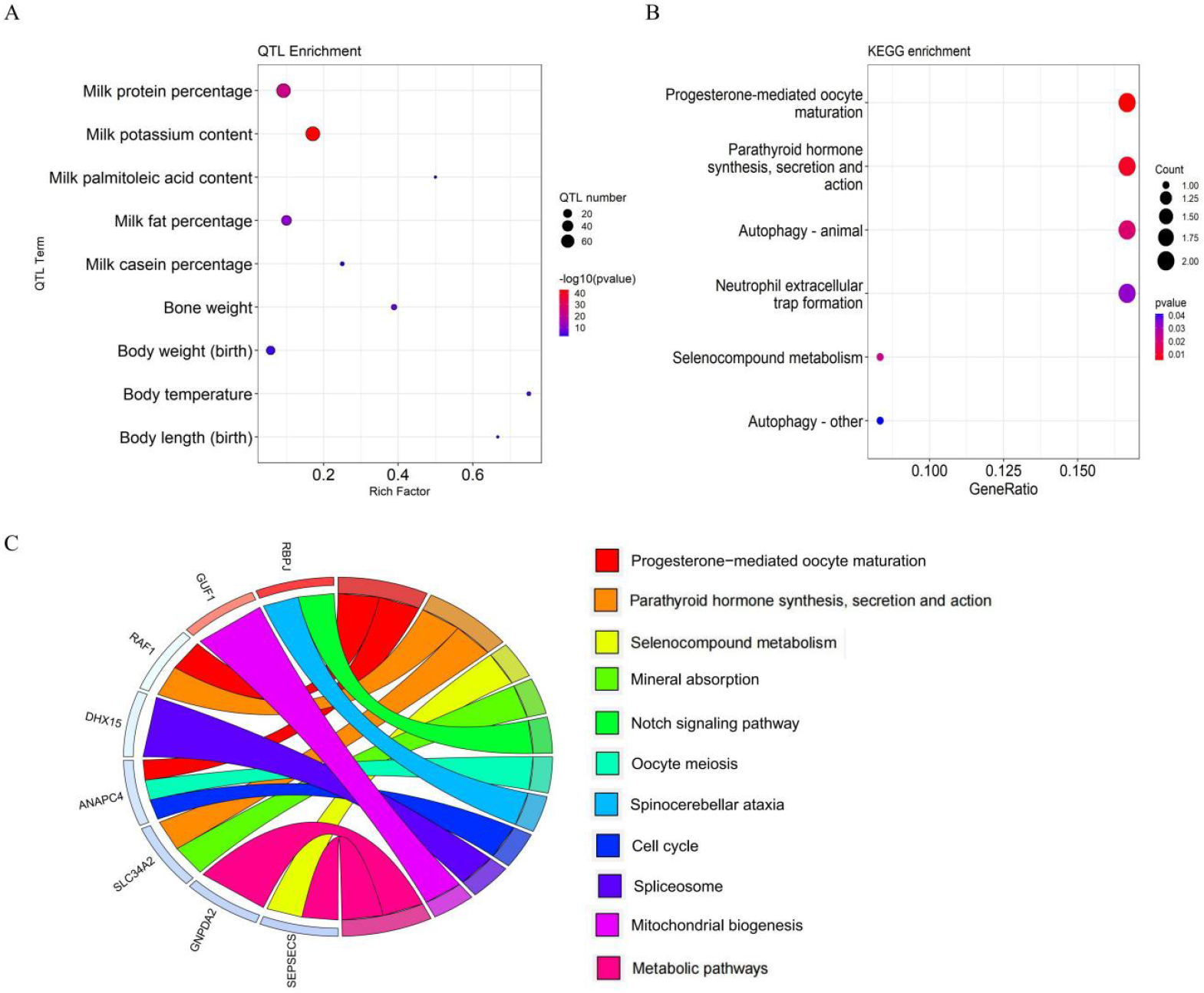
Functional enrichment of candidate genes and QTLs. (A) The enriched traits identified in QTL enrichment. The area of the bubbles represents the number of observed QTLs for that class, while the color represents the -log10(p) value scale. The X-axis shows the richness factor for each QTL, representing the ratio of number of QTLs and the expected number of that QTL. (B) KEGG pathway enrichment of the candidate genes. The size of each bubble represents the number of genes enriched in the pathway, and the color intensity indicates the statistical significance. The X-axis represents the enrichment score, while the Y-axis shows the KEGG pathway terms. (C) The circles show the pathways in which candidate genes are involved.

Candidate Region 1 on BTA6 contained 9 protein coding genes and 3 non-coding genes, Region 2 on BTA6 contained one protein coding gene *KCTD8*, and Region 3 on BTA18 contained one protein coding gene *MAF*. Functional enrichment analysis of candidate genes showed that they were mainly enriched in progesterone-mediated oocyte maturation, parathyroid hormone synthesis, secretion and action, and selenocompound metabolism signaling pathways (Fig. 3B). Specifically, the candidate gene *ANAPC4* was enriched in progesterone-mediated oocyte maturation and participated in the process of oocyte meiosis; *SEPSECS* was enriched in selenocompound metabolism; *RBPJ* was involved in Notch signaling pathway, which played a vital role during growth and development (Fig. 3C).

### Colocalization Analysis of the Candidate Variants

Traditionally, the nearest genes to lead SNPs were recognized as risk genes at GWAS loci. However, multiple lead SNPs and distinct candidates might be reported within the same locus. In our candidate Region 1, 45 significant SNPs clustered into 4 peaks all showed a relatively high positive effect (0.45-0.85) for AFS (Fig. 2C), indicating the possibility of more than one causal gene in the region. Rs136363104 was identified as a cis-eQTL that could regulate the expression of *ANAPC4* in liver, and as cis-sQTL regulating its splicing in lymph node and blood (Fig. 4; Table S4, Additional file 1). Considering the previous functional enrichment of *ANAPC4* and its role in progesterone-mediated oocyte maturation, it suggests that *ANAPC4* and rs136363104 could be the putative causal gene and variant for puberty. Meanwhile, rs208065122, a missense mutation identified in *ZCCHC4*, had a strong linkage (R^2^ > 0.8) with rs136363104 and the lead SNP rs211544714, indicating a potential impact on AFS.

**Fig. 4.**
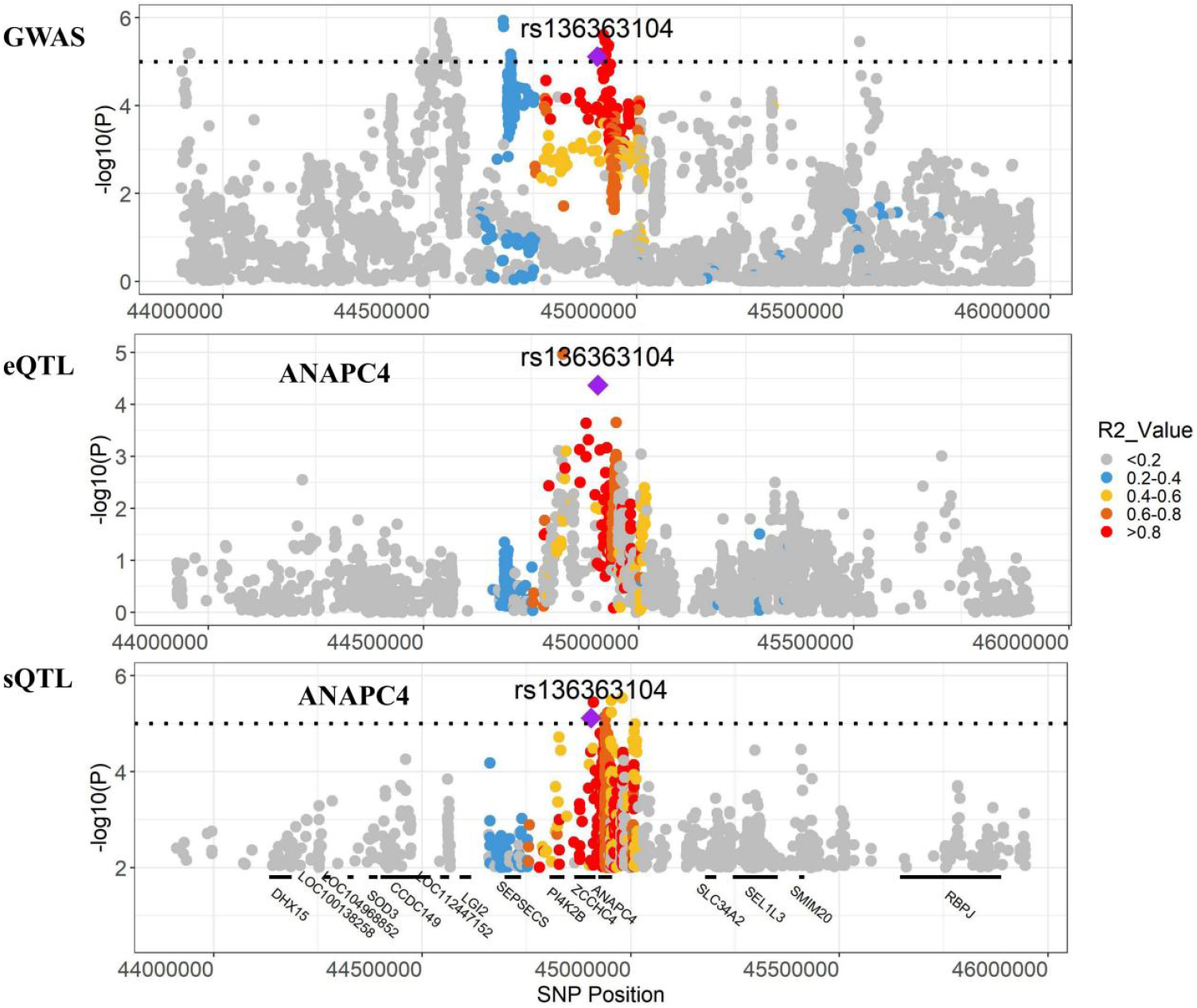
Colocalization analysis of GWAS, eQTL, and sQTLs in the candidate region. Locus zoom plot of the GWAS association (upper), eQTL for the association between ANAPC4 expression and genotype in liver (middle), and sQTL between ANAPC4 splicing and genotype in blood (bottom). The purple diamonds represent the putative causal variant (rs136363104) of ANAPC4. The color indicated on the right panel represents the linkage disequilibrium R^2^ value between each SNP and the inferred causal SNP rs136363104.

Additionally, eight cis-sQTLs located at BTA6:44809081-45007127 of Region 1 were detected to regulate the alternative splicing of the coding genes *RBPJ, SEPSECS* and *SLC34A2*, as well as three non-coding RNAs (Table S4, Additional file 1). Within the interval BTA6:44369507-44569075 of Region 1, the splicing of *CCDC149, DHX15, RACK1, SEPSECS, ZCCHC4*, and LOC112447057 was regulated by 10 cis-sQTLs. Rs135085994, rs135261840, and rs132774279 were three cis-eQTLs regulating the expression of *SOD3*. In Region 2, one cis-eQTLs, 12 cis-sQTLs, and two trans-sQTLs were identified, regulating *GNPDA2, GUF1, KCTD8*, and LOC104968873 (Table S4, Additional file 1). Among them, rs42432508 and rs42432506 could regulate the splicing of *GUF1* in ovary. Although no sQTL or eQTL was identified in candidate Region 3, previously reported QTLs in this region were associated with Anti-Mullerian Hormone (AMH), a dimeric glycoprotein secreted by granulosa cells of female follicles that inhibits the recruitment and growth of follicles and regulates the differentiation and development of reproductive ducts. A high level of AMH indicates a larger oocytes stock.

### Expression Validation of Candidate Genes in Blood RNA-seq Data

Two groups of samples (20 individuals each) with significant differences in both AFS and GEBVs were used to explore the differences in gene expression (Fig. 5A). There were also significant differences between the high and low AFS group for RNA-seq counts based on principle component analysis (Fig. 5B). The analysis of the differently expressed genes (DEGs) were detected using the weighted least square method based on a linear model, with multiple tests corrected using Bayesian methods. A total of 1740 significantly up-regulated genes and 1777 down-regulated genes were found in the group with low AFS compared to high group (Fig. 5C; Table S5, Additional file 1). Five candidate genes identified based on GWAS were also DEGs. Of these, four significant genes (*ANAPC4, DHX15, GUF1*, and *SEPSECS*) were up-regulated, while the other one (*SOD3*) was down-regulated.

**Fig. 5.**
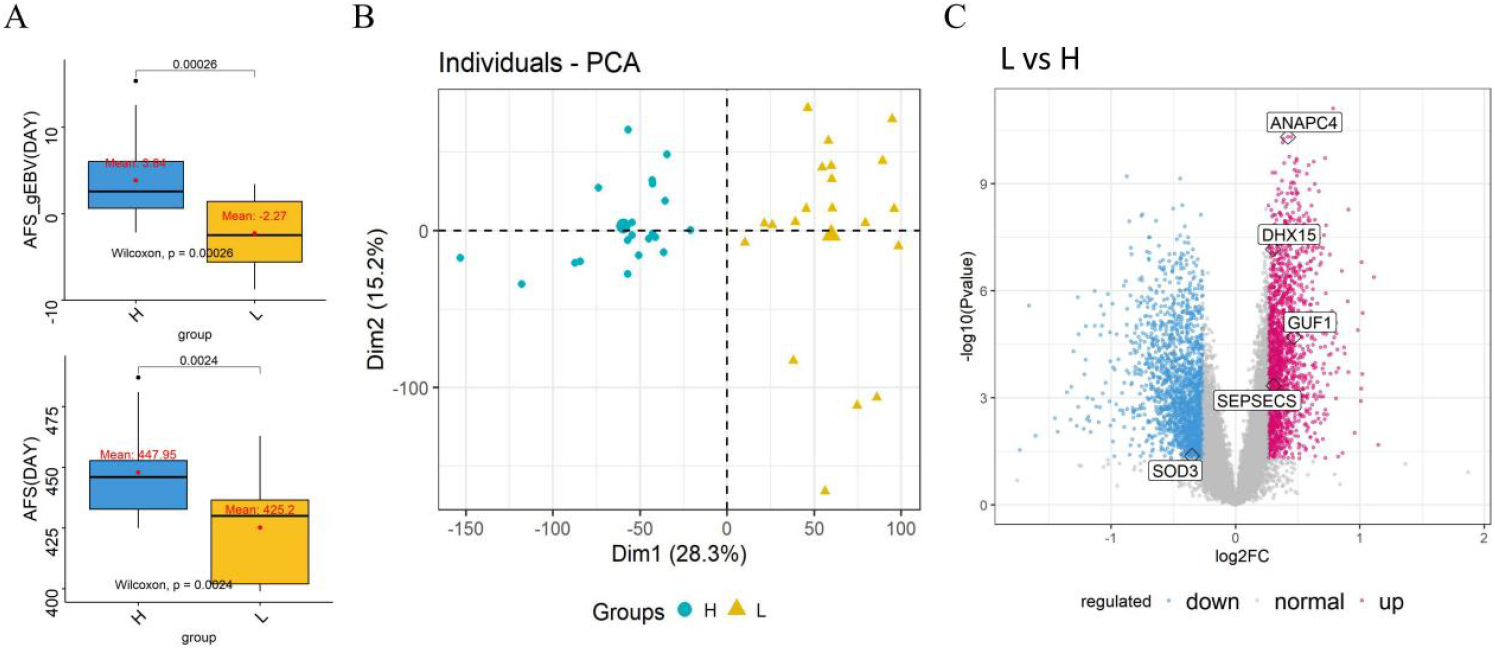
Sample Clustering and gene differential expression analysis between high and low AFS groups. (A) Box plot displaying the significant differences of phenotype and GEBV in high and low groups. (B) Principle component analysis for samples on the basis of gene expression. (C) Volcano plot showing significantly expressed genes. The light blue and pink dots denote significantly down-regulated and up-regulated genes, respectively. H and L represent high and low AFS groups, respectively.

Colocalization indicated that rs136363104 was a putative causal variant for AFS. Further analysis showed the expression of ANAPC4 in the GG haplotype of rs136363104 was significantly higher than that in the AA haplotype in blood (Fig. 6A). Transcript analysis indicated that the GG haplotype showed high expression of ENSBTAT00000000852.6, while the expression level of ENSBTAT00000078795.1 was consistently low across all haplotypes.

**Fig. 6.**
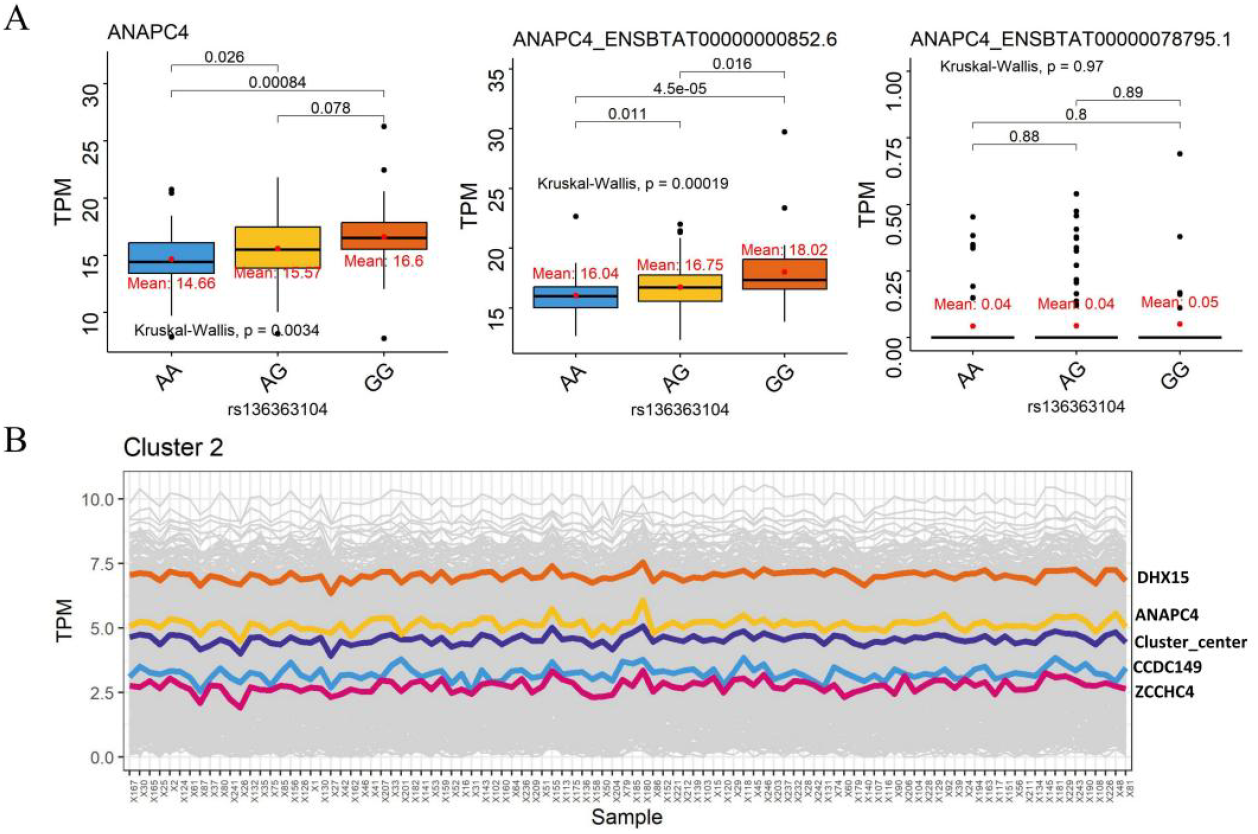
The expression validation of *ANAPC4* in blood. (A) Box plots shows the differences in the expression of candidate gene *ANAPC4*, and its two transcripts ENSBTAT00000000852.6 and ENSBTAT00000078795.1 between three haplotypes (rs136363104). The mean and pairwise comparisons p-value of each group were marked. (B) Cluster 2 of genes by k-means clustering. Cluster center and four candidate genes was displayed by different colors.

To understand the expression patterns and their co-regulation relationships of the candidate genes, those not expressed in 50% of the samples were removed, resulting in a final set of 15,571 genes. K-means clustering divided the genes into 20 clusters, ranging from 444 genes (Cluster 3) to 1298 genes (Cluster 19). The 13 candidate genes were distributed across 7 clusters (Fig. S6, Additional file 2). Notably, *ANAPC4* exhibited similar expression patterns with *DHX15, CCDC149*, and *ZCCHC4* in Cluster 2 (Fig. 6B), suggesting potential synergistic regulatory functions in growth, development and sexual maturation.

In summary, rs136363104 was proposed as the putative causal variant for AFS in the Chinese Holstein population, primarily through its regulation of *ANAPC4* expression. Higher *ANAPC4* expression levels might advance the puberty and reduce AFS. However, due to the complexity of AFS, its regulation likely involves polygenes with subtle individual variation effects.

## DISCUSSION

### Genetic Parameters Analysis

The average AFS was 14.0 months (422 days) while AFC was 23.8 months (716 days) in our study, comparable to the international average level (Hutchison et al., 2017; Heise et al., 2018; Kgari et al., 2022b). A previous study suggested the optimal AFC to maximize production in Holstein cows was 21 months (Hutchison et al., 2017), indicating potential for further improvement in Chinese Holsteins.

The heritability for AFS estimated in this study was moderate (0.276±0.025), comparable to estimates from previous studies, such as 0.25 in Dutch Friesians (JANSEN et al., 1987), 0.299 in Jersey × Red Sindhi crossbred cows (Vinothraj et al., 2016), and 0.22 in Ayrshire and Friesian cows (Mäntysaari et al., 2002). However, low heritability estimates for AFS has been reported, such as 0.107 in Iranian Holstein heifers (Eghbalsaied, 2011), 0.12 and 0.13 in Canadian Holsteins (Raheja et al., 1989; Jamrozik et al., 2005), and as low as 0.02 in South African Holstein heifers (Kgari et al., 2022a). High heritability estimates were also observed, with a value of 0.58 in Chinese Holsteins from a herd in Ningxia (Hu et al., 2023). Aside from AFS, the other six fertility traits all had low heritability, ranging from 0.001 to 0.131. Notably, the heritability of AFC (0.057) in our study was slightly lower than the 0.08 in South African Holstein heifers (Kgari et al., 2022a), 0.13 in multiple breeds of New Zealand (Grosshans et al., 1997), 0.22 in Jersey × Red Sindhi crossbred cows (Vinothraj et al., 2016), but higher than the 0.027 reported in Iranian Holstein heifers (Eghbalsaied, 2011).

The discrepancy in heritability estimates might be due to the environmental and management differences, which have been shown to significantly affect fertility traits (Slagboom et al., 2021; Mancin et al., 2024). Additionally, several factors can influence the estimation of heritability, including the genetic variation of varied populations (Notter, 1999), different statistical models for analysis (Yadav et al., 2023), the interaction of breeds with environmental conditions, and the reliability, size, and quality of the data.

The genetic correlations between AFS and AFC were high and favorable (0.93 ± 0.03). This was consistent with previous studies, which reported correlations of 0.91 in South African Holsteins (Kgari et al., 2022a), 0.98 in Polish Holsteins (Jagusiak et al., 2006), 0.99 in Czech Holsteins (Brzáková et al., 2019), and approximately 1.0 in Thailand up-graded Holsteins (Buaban et al., 2015), respectively. These results indicates that the two traits likely share the same physiological and genetic basis, and the improvement of AFC is heavily dependent on AFS. Based on these results, we analyzed the genetic regions and variations of AFS with relatively high heritability to explore AFC, which has low heritability in this study.

### Candidate Regions Analysis

To avoid the adverse effects of premature breeding, calves must reach both sexual and body maturation to achieve the optimal calving state. It has been reported that the first insemination is recommended when German Holsteins have reached a weight above 370 kg (average 406.46 ± 32.90 kg) (Yin et al., 2018). In Israel, heifers are generally inseminated at the age of 14 months, with a weight above 350 kg, and a withers height above 125 cm (Weller et al., 2022a). For Chinese Holstein heifers in this project, the insemination criteria were a weight greater than 380 kg, an age greater than 400 days and combined with estrus monitoring. Thus, AFS is a composite trait that included both growth and development (body weight), and sexual maturity (estrus).

In this study, three candidate regions were identified for AFS, encompassing five 200 kb windows. Particularly, one region on BTA6:43688837-45007127, containing three adjacent windows within 1.32 Mb, was rich in genes and QTLs. Previous study has been reported that the region from 25 to 53 Mbp on BTA6 contains a significant concentration of SNPs strongly associated with production and growth traits, including QTLs for birth weight (Casas et al., 2000; Kneeland et al., 2004; Gutiérrez-Gil et al., 2009), yearling weight (Casas et al., 2000), and pre- and post-weaning body weight gain (Kneeland et al., 2004). A GWAS study of a large size population, including 27,707 *Bos indicus, Bos taurus* and crossbred cattle, revealed that this region is also highly associated with heifer puberty (Forutan et al., 2024). The region explained 2.3% genetic variance of AFS in our study. Consistent with previous results, candidate Region 1 (43.6 to 45.0 Mbp on BAT6), with the largest variance explanation, contained 14 QTLs for birth weight, 5 QTLs for body weight gain, 2 QTLs for birth body length, and 1 for yearling weight (Snelling et al., 2010; Gutiérrez Gil et al., 2012; Lu et al., 2013). A large number of QTLs in this region were also found to be associated with milk trait. For instance, there were 74 QTLs associated with milk protein percentage (Buitenhuis et al., 2016; Olsen et al., 2016), 32 QTLs with milk fat percentage (Olsen et al., 2016), 77 QTLs with milk potassium content (Buitenhuis et al., 2015), 5 QTLs with milk casein percentage/content (Buitenhuis et al., 2016), and 3 QTLs with milk yield (Viale et al., 2017; Weikard et al., 2012). Candidate Region 1 also incorporated QTLs related to reproduction, such as conception rate (Parker Gaddis et al., 2016), calving to conception interval (Müller et al., 2017), inseminations per conception (Höglund et al., 2015), non-return rate (Höglund et al., 2015), and calving ease (Cole et al., 2011).

Similarly, results showed that candidate Region 2 also comprised QTLs related to production (birth weight, weaning weight, yearling weight, and body weight gain) (Snelling et al., 2010), milk (milk kappa-casein percentage, milk glycosylated kappa-casein percentage, and milk unglycosylated kappa-casein percentage) (Buitenhuis et al., 2016), health (bovine tuberculosis susceptibility) (Richardson et al., 2016), and exterior (udder swelling score) (Michenet & Barbat et al., 2016; Michenet & Saintilan et al., 2016). In candidate Region 3, three QTLs were reported: one associated with AMH (Gobikrushanth et al., 2018), and two with milk caproic/caprylic content (Ibeagha-Awemu et al., 2016). It indicated this region might regulate reproductive duct differentiation and development, increasing oocyte capacity by influencing AMH levels. In summary, the analysis confirmed the pleiotropy of the candidate regions, especially the comprehensive regulation in production, milk, and reproduction traits.

### Putative Causal Gene analysis

Combining positional information with eQTL and sQTL results, 12 coding genes (*ANAPC4, RBPJ, SEPSECS, SLC34A2, ZCCHC4*, CCDC149, *GNPDA2, GUF1, DHX15, SOD3, GUF1* and *GNPDA2*) were identified as key candidate genes for AFS. In humans, *ZCCHC4, RBPJ, SEPSECS* have been found to be associated with body weight index (BMI) (Snelling et al., 2010; Schoeler et al., 2023), *RBPJ* with waist-to-hip ratio adjusted for BMI (Pulit et al., 2019), and *DHX15* and *SLC34A2* with height (Yengo et al., 2022). Consistent with these results, missense mutations in *ZCCHC4* (rs208065122), *SEPSECS* (rs381489766), and a synonymous mutation in *ANAPC4* (rs132745273) were identified as significant SNPs for carcass weight and eye muscle area in Korean Hanwoo cattle (Bhuiyan et al., 2018).

As AFS is a composite trait, in addition to candidate genes regulating production (mainly body weight), a key finding was our proposed of *ANAPC4* as the putative causal gene for puberty in Holstein heifers. Mammalian oocytes are arrested at prophase I before puberty until luteinizing hormone (LH) induces resumption of meiosis of follicle-enclosed oocytes. ANAPC4 (Anaphase Promoting Complex Subunit 4) is a component of the anaphase promoting complex/cyclosome (APC/C), an E3 ubiquitin ligase that controls the progression through mitosis and meiosis of the cell cycle, playing a crucial role in oocyte maturation and early embryonic development. (Malhotra et al., 2016). Variants in *ANAPC4* might influence the timing of puberty onset and first estrus. Previous study has shown that variants in *ANAPC4* was associated with age at menarche (P = 4.65×10^−8^) (Kichaev et al., 2019) and age at first sexual intercourse (Mills et al., 2021) in humans. Herein, a putative causal variant (rs136363104) for puberty was identified by regulating the expression of *ANAPC4* in blood. However, it should also be highlighted that gene expression in blood might only help to explain some of the mutations affecting AFS. Future studies in uterus and ovary are recommended, as several genes responsible for development and reproduction are expressed at much higher levels.

Additionally, *RBPJ* was a candidate gene significantly associated with AFS suggested by gene-based GWAS analysis (Fig. S7, Additional file 2). RBPJ (Recombination Signal Binding Protein For Immunoglobulin Kappa J Region) is an important transcriptional regulator in the Notch signaling pathway, involved in cell-cell communication that regulates a broad spectrum of cell-fate determinations during individual development (Friedrich et al., 2022). Altered RBPJ expression or function could impact mammary gland maturation and reproduction. RBPJ could confer on-time uterine lumen shape transformation via physically interacting with uterine estrogen receptor (ERα), which is essential for mouse uterine closure and correct alignment of implanted embryos (Zhang et al., 2014).

## CONCLUSIONS

In this study, we identified three genomic regions associated with AFS that explained a total of 0.91% genetic variance in Chinese Holstein heifers. The candidate genes were involved in growth, development and sexual maturity. Combining GWAS with eQTL and sQTL analysis, we identified *ANAPC4, RBPJ, SEPSECS, SLC34A2, ZCCHC4*, CCDC149, *GNPDA2, GUF1, DHX15, SOD3, GUF1* and *GNPDA2* as candidate genes for AFS. Specifically, *ANAPC4* and rs136363104 were proposed as the putative causal gene and variant for puberty, respectively. Our results suggest that the regions on BTA6 regulating AFS may have pleiotropic and functionally correlated effects with several traits related to growth, development and puberty. It also indicates that numerous candidate variations affect a set of genes, collectively explaining the genetic variance for AFS. In summary, considering the moderate heritability of AFS and its high genetic correlations with AFC, our findings suggested that there is potential to reduce rearing costs and maximize production efficiency by genetic selection for AFS in Chinese Holstein population.

## Abbreviations

AFS: age at first service
AFC: age at first calving
QTL: quantitative trait loci
eQTL: expression quantitative trait locus
sQTL: splicing quantitative trait locus
GWAS: genome-wide association study
GEBV: genomic estimated breeding value
AGEP: age at puberty
RNA-seq: RNA sequencing
LD: linkage disequilibrium
GLM: general liner model
CV: coefficient of variation
IFLh: interval from first to last inseminations of heifers
CRh: conception rate of heifers
NS: number of repeated service
GL: gestation length
BiWh: birth weight of heifers

## Supplementary Information

**Additional file 1** (https://figshare.com/s/da6c40422c67302dbe6d):

**Table S1**. The values of *Me* and significance threshold for all chromosomes.

**Table S2**. The windows (200kb) of top 1% variance explaination.

**Table S3**. The significant SNP identified by GWAS.

**Table S4**. The colocalization of significant SNPs with eQTL or sQTL in the candidate region.

**Table S5**. The differentially expressed genes□:DEGs□: between low and high AFS groups in blood.

**Additional file 2** (https://figshare.com/s/4a94a5a91d6cf49847a9):

**Fig. S1**. SNP density distribution after qulity control.

**Fig. S2**. The quantile-quantile plots of GWAS results.

**Fig. S3**. Variants analysis of the candidate Region 1.

**Fig. S4**. The frequency of each QTL class in the candidate regions.

**Fig. S5**. The composition of the Milk (A), Production (B), and Reproduction(C) QTLs class.

**Fig. S6**. K-means cluster of the genes in blood.

**Fig. S7**. Gene-based GWAS analysis by MAGMA.

## Notes

### Ethics approval and consent to participate

Not applicable

### Consent for publication

Not applicable.

### Availability of data and material

The datasets generated during the current study are not publicly available, but are available from the corresponding author on reasonable request.

### Authors’ contributions

Conceived and designed the research: Yu Jiang; Analyzed the data: Cuili Pan, Longgang Ma, Zhuangbiao Zhang, Ao Wang, Lulu Wang, Fengting Bai; Sample and data collection: Lulu Wang, Guanglei Liu, Kai Zhu, Xinzhe Lv; Wrote the manuscript: Cuili Pan; Modified the manuscript: Yu Jiang, Xihong Wang, Yudong Cai, Yun Ma, Yachun Wang. All authors read and approved the final manuscript.

## Acknowledgments

We thank the high-performance computing platform of Northwest A&F University for providing the computing resources.

